# Persistent cross-species transmission systems dominate Shiga toxin-producing *Escherichia coli* O157:H7 epidemiology in a high incidence region: a genomic epidemiology study

**DOI:** 10.1101/2024.04.05.588308

**Authors:** Gillian A.M. Tarr, Linda Chui, Kim Stanford, Emmanuel W. Bumunang, Rahat Zaheer, Vincent Li, Stephen B. Freedman, Chad R. Laing, Tim A. McAllister

**Affiliations:** Division of Environmental Health Sciences, School of Public Health, University of Minnesota, MMC807, Room 1240 Mayo, 420 Delaware Street SE, Minneapolis, MN, 55455, USA; Alberta Precision Laboratories, Alberta Public Health, Room 1B2.19 Walter Mackenzie Health Sciences Centre, 8440-112th Street, Edmonton, Alberta, T6G 2J2, Canada; Department of Laboratory Medicine and Pathology, University of Alberta, Edmonton, Alberta, Canada; Department of Biological Sciences, University of Lethbridge, 4401 University Drive, Lethbridge, Alberta, T1K 3M4, Canada; Agriculture and Agri-Food Canada, Lethbridge Research and Development Centre, 5403 – 1 Avenue South. P.O. Box 3000, Lethbridge, Alberta, T1J 4B1, Canada; Sections of Pediatric Emergency Medicine and Gastroenterology, Department of Pediatrics, Alberta Children’s Hospital and Alberta Children’s Hospital Research Institute, Cumming School of Medicine, University of Calgary, C4-634 28 Oki Drive NW, Calgary, Alberta, T3B 6A8, Canada; National Center for Animal Diseases Lethbridge Laboratory, Canadian Food Inspection Agency, 225090 Township Road 91, Lethbridge County, Alberta, T1J 5R7, Canada

**Keywords:** *E. coli* O157, STEC, genomic epidemiology, zoonosis

## Abstract

**Background:** Several areas of the world suffer notably high incidence of Shiga toxin-producing *Escherichia coli*, among them Alberta, Canada. We assessed the impact of persistent cross-species transmission systems on the epidemiology of *E. coli* O157:H7 in Alberta.

**Methods:** We sequenced and assembled 229 *E. coli* O157:H7 isolates originating from collocated cattle (n=108) and human (n=121) populations from 2007-2015 in Alberta. We constructed a timed phylogeny using BEAST2 using a structured coalescent model. We then extended the tree with human isolates through 2019 (n=430) to assess the long-term disease impact of locally persistent lineages. Shiga toxin gene (*stx*) profile was determined for all isolates.

**Results:** During 2007 to 2015, we estimated 108 (95% HPD 104, 112) human lineages arose from cattle lineages, and 14 (95% HPD 5, 23) from other human lineages; i.e., 88.5% of human lineages arose from cattle lineages. We identified 11 persistent lineages local to Alberta, which were associated with 38.0% (95% CI 29.3%, 47.3%) of human isolates. Of 117 isolates in locally persistent lineages, 6.0% carried only the Shiga toxin gene *stx2a* and the rest both *stx1a* and *stx2a*. During the later period, six locally persistent lineages continued to be associated with human illness, including 74.7% (95% CI 68.3%, 80.3%) of reported cases in 2018 and 2019. The *stx* profile of isolates in locally persistent lineages shifted from the earlier period, with 51.2% encoding only *stx2a*.

**Conclusions:** Our study identified multiple locally evolving lineages transmitted between cattle and humans persistently associated with *E. coli* O157:H7 illnesses for up to 13 years. Of concern, there was a dramatic shift in locally persistent lineages toward strains with the more virulent *stx2a*-only profile. Locally persistent lineages may be a principal cause of the high incidence of *E. coli* O157:H7 in locations such as Alberta and offer opportunities for understanding the disease ecology supporting *E. coli* O157:H7 persistence, as well as for local prevention efforts.

## Introduction

Several areas around the globe experience exceptionally high incidence of Shiga toxin-producing *Escherichia coli* (STEC), including the virulent serotype *E. coli* O157:H7. These include Scotland,^1^ Ireland,^2^ Argentina,^3^ and the Canadian province of Alberta.^4^ All are home to large populations of agricultural ruminants, STEC’s primary reservoir. However, there are many regions with similar ruminant populations where STEC incidence is unremarkable. What differentiates high risk regions is unclear. Moreover, with systematic STEC surveillance only conducted in limited parts of the world,^5^ there may be unidentified regions with exceptionally high disease burden.

STEC infections can arise from local reservoirs, transmitted through food, water, direct animal contact, or contact with contaminated environmental matrices. The most common reservoirs include domesticated ruminants such as cattle, sheep, and goats. Animal contact and consumption of contaminated meat and dairy products are significant risk factors for STEC, as are consumption of leafy greens, tomatoes, and herbs and recreational swimming that have been contaminated by feces from domestic ruminants.^6^ While STEC has been isolated from a variety of other animal species and outbreaks have been linked to species such as deer^7^ and swine,^8^ it is unclear what roles they play as maintenance or intermediate hosts. STEC infections can be imported through food items traded nationally and internationally, as has been seen with *E. coli* O157:H7 outbreaks in romaine lettuce from the United States.^9^ Secondary transmission is believed to cause approximately 15% of cases, but transmission of the pathogen is not believed to be sustained through person-to-person transmission over the long term.^10,11^

The mix of STEC infection sources in a region directly influences public health measures needed to control disease burden. Living near cattle and other domesticated ruminants has been linked to STEC incidence, particularly for *E. coli* O157:H7.^2,12–16^ These studies suggest an important role for local reservoirs in STEC epidemiology. A comprehensive understanding of STEC’s disease ecology would enable more effective investigations into potential local transmission systems and ultimately their control. Here, we take a phylodynamic, genomic epidemiology approach to more precisely discern the role of the cattle reservoir in the dynamics of *E. coli* O157:H7 human infections. We focus on the high incidence region of Alberta, Canada to provide insight into characteristics that make the pathogen particularly prominent in such regions.

## Methods

### Study Design and Population

We conducted a multi-host genomic epidemiology study in Alberta, Canada. Our primary analysis focused on 2007 to 2015 due to the availability of isolates from intensive provincial cattle studies.^17–20^ These studies rectally sampled feces from individual animals, hide swabs, fecal pats from the floors of pens of commercial feedlot cattle, or feces from the floors of transport trailers. In studies of pens of cattle, samples were collected from the same cattle at least twice over a four to six-month period. A one-time composite sample was collected from cattle in transport trailers, which originated from feedlots or auction markets in Alberta. To select both cattle and human isolates, we block randomized by year to ensure representation across the period. We define isolates as single bacterial species obtained from culture. We sampled 123 *E. coli* O157 cattle isolates from 4,660 available. Selected cattle isolates represented 7 of 12 cattle study sites and 56 of 89 sampling occasions from the source studies.^17–20^ We sampled 123 of 1,148 *E. coli* O157 isolates collected from cases reported to the provincial health authority (Alberta Health) during the corresponding time period (Supplemental Information).

In addition to the 246 isolates for the primary analysis, we contextualized our findings with two additional sets of *E. coli* O157:H7 isolates (Figure 1): 445 from Alberta Health from 2009 to 2019 and already sequenced as part of other public health activities and 1,970 from the U.S. and elsewhere around the world between 1999 and 2019. The additional Alberta Health isolates were sequenced by the National Microbiology Laboratory (NML)-Public Health Agency of Canada (Winnipeg, Manitoba, Canada) as part of PulseNet Canada activities. Isolates sequenced by the NML for 2018 and 2019 constituted the majority of reported *E. coli* O157:H7 cases for those years (217 of 247; 87.9%). U.S. and global isolates from both cattle and humans were identified from previous literature (n=104)^21^ and BV-BRC (n=193). As both processed beef and live cattle are frequently imported into Alberta from the U.S., we selected additional *E. coli* O157:H7 sequences available through the U.S. CDC’s PulseNet BioProject PRJNA218110. From 2010-2019, 6,791 O157:H7 whole genome sequences were available from the U.S. PulseNet project, 1,673 (25%) of which we randomly selected for assembly and clade typing.

**Figure 1.**
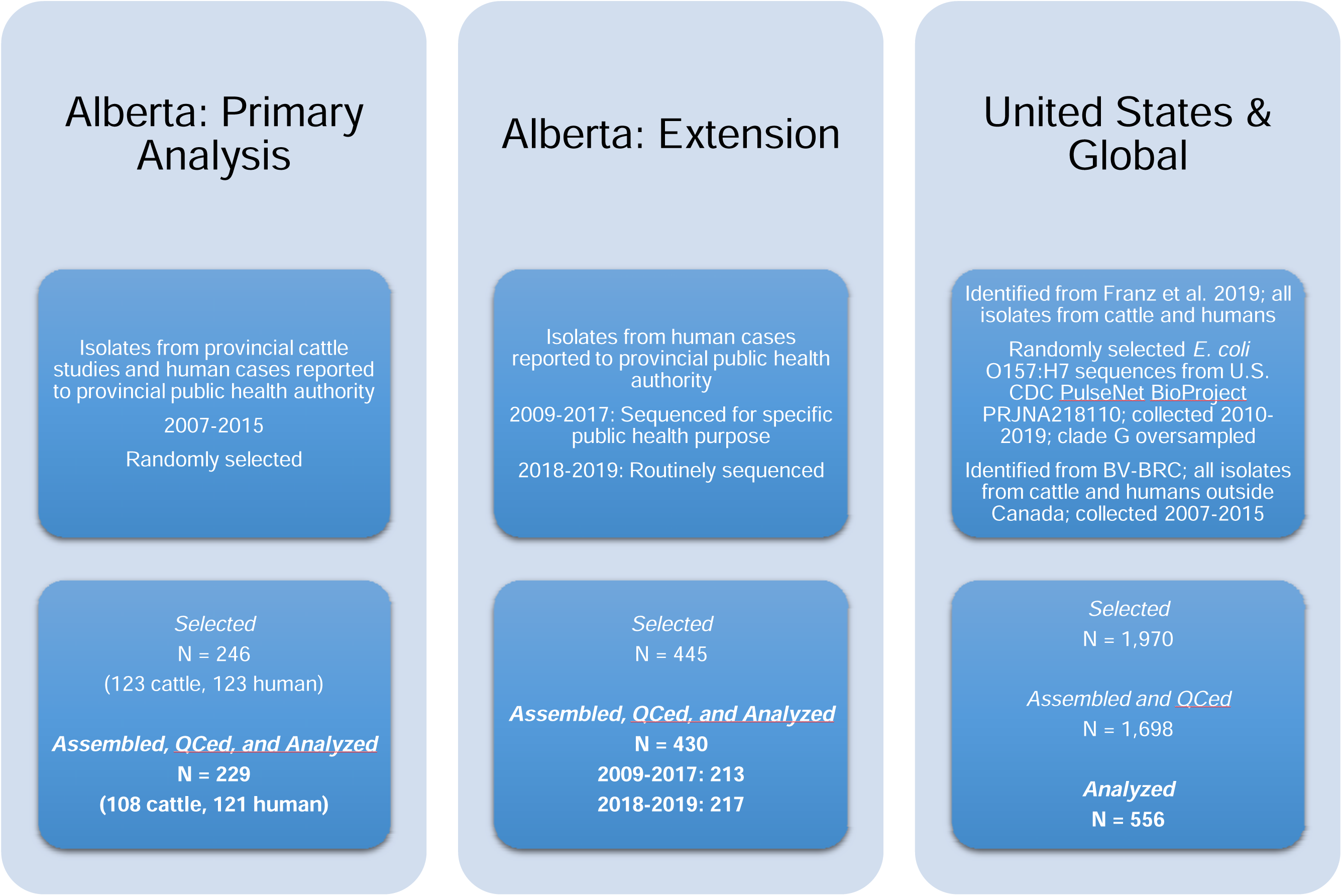
*E. coli* O157:H7 isolates selected for the study and assembled. Three sets of isolates, all originating from cattle or humans, were included in the study.

This study was approved by the University of Calgary Conjoint Health Research Ethics Board, #REB19-0510. A waiver of consent was granted, and all case data were deidentified.

### Whole Genome Sequencing, Assembly, and Initial Phylogeny

The 246 isolates for the primary analysis were sequenced using Illumina NovaSeq 6000 and assembled into contigs using the Unicycler v04.9 pipeline, as described previously (BioProject PRJNA870153).^22^ Raw read FASTQ files were obtained from Alberta Health for the additional 445 isolates sequenced by the NML and from NCBI for the 152 U.S. and 54 global sequences. We used the SRA Toolkit v3.0.0 to download sequences for U.S. and global isolates using their BioSample (i.e. SAMN) numbers. The corresponding FASTQ files could not be obtained for 6 U.S. and 7 global isolates we had selected (Figure 1).

PopPUNK v2.5.0 was used to cluster Alberta isolates and identify any outside the O157:H7 genomic cluster (Supplemental Figure S1).^23^ For assembling and quality checking (QC) all sequences, we used the Bactopia v3.0.0 pipeline.^24^ This pipeline performed an initial QC step on the reads using FastQC v0.12.1, which evaluated read count, sequence coverage, and sequence depth, with failed reads excluded from subsequent assembly. None of the isolates were eliminated during this step for low read quality. We used the Shovill v1.1.0 assembler within the Bactopia pipeline to *de novo* assemble the Unicycler contigs for the primary analysis and raw reads from the supplementary datasets. Trimmomatic was run as part of Shovill to trim adapters and read ends with quality lower than 6 and discard reads after trimming with overall quality scores lower than 10. Bactopia generated a quality report on the assemblies, which we assessed based on number of contigs (<500), genome size (≥5.1 Mb), N50 (>30,000), and L50 (≤50).

Low-quality assemblies were removed. This included 1 U.S. sequence, for which 2 FASTQ files had been attached to a single BioSample identifier; the other sequence for the isolate passed all quality checks and remained in the analysis. Additionally, 16 sequences from the primary analysis dataset and 4 from the extended Alberta data had a total length <5.1 Mb. These sequences corresponded exactly to those identified by the PopPUNK analysis to be outside the primary *E. coli* O157:H7 genomic cluster (Supplemental Figure S1). Finally, although all isolates were believed to be of cattle or clinical origin during initial selection, detailed metadata review identified 1 isolate of environmental origin in the primary analysis dataset and 8 that had been isolated from food items in the extended Alberta data. These were excluded. We used STECFinder v1.1.0^25^ to determine Shiga toxin gene (*stx*) profile and confirm the *E. coli* O157:H7 serotype using the *wzy* or *wzx* O157 O-antigen genes and detection of the H7 H-antigen.

Bactopia’s Snippy workflow, which incorporates Snippy v4.6.0, Gubbins v3.3.0, and IQTree v2.2.2.7, followed by SNP-Sites v2.5.1, were used to generate a core genome SNP alignment with recombinant blocks removed. The maximum likelihood phylogeny of the core genome SNP alignment generated by IQTree was visualized in Microreact v251. The number of core SNPs between isolates was calculated using PairSNP v0.3.1. Clade was determined based on the presence of at least one defining SNP for the clade as published previously.^26^ Isolates were identified to the clade level, except for clade G where we separated out subclade G(vi).

After processing, we had 229 isolates (121 human, 108 cattle) in our primary sample and 430 additional Alberta Health isolates (Figure 1, Supplemental Data File). We had 178 U.S. or global isolates from previous literature (n=88; U.S. n=41, global n=47) and BV-BRC (n=90; U.S. n=75, global n=15). Of the 1,673 isolates randomly sampled from the U.S. PulseNet project, 1,560 were successfully assembled and passed QC. These included 309 clade G isolates, all of which we included in the analysis; we also randomly sampled and included 69 non-clade G isolates from this sample.

### Phylodynamic and Statistical Analyses

For our primary analysis, we created a timed phylogeny, a phylogenetic tree on the scale of time, in BEAST2 v2.6.7 using the structured coalescent model in the Mascot v3.0.0 package with demes for cattle and humans (Supplemental Table S1). Sequences were down-sampled prior to analysis if within 0-2 SNPs and <3 months from another sequence from the same host type, leaving 115 human and 84 cattle isolates in the primary analysis (Supplemental Table S1). The analysis was run using four different seeds to confirm that all converged to the same solution, and tree files were combined before generating a maximum clade credibility (MCC) tree. State transitions between cattle and human isolates over the entirety of the tree, with their 95% highest posterior density (HPD) intervals, were also calculated from the combined tree files. We determined the influence of the prior assumptions on the analysis (Supplemental Table S1) with a run that sampled from the prior distribution (Supplemental Information). We conducted a sensitivity analysis in which we randomly subsampled 84 of the human isolates so that both species had the same number of isolates in the analysis.

Locally persistent lineages (LPLs) were identified based on following criteria: 1) a single lineage of the MCC tree with a most recent common ancestor (MRCA) with ≥95% posterior probability; 2) all isolates ≤30 core SNPs from one another; 3) contained at least 1 cattle isolate; 4) contained ≥5 isolates; and 5) the isolates were collected at sampling events (for cattle) or reported (for humans) over a period of at least 1 year. We counted the number of isolates associated with LPLs, including those down-sampled prior to the phylodynamic analysis. We conducted sensitivity analyses examining different SNP thresholds for the LPL definition.

From non-LPL isolates, we estimated the number of local transient isolates vs. imported isolates. For the 121 human *E. coli* O157:H7 isolates in the primary sample prior to down-sampling, we determined what portion belonged to locally persistent lineages and what portion were likely to be from local transient *E. coli* O157:H7 populations vs. imported. Human isolates within the LPLs were enumerated (n=46). The 75 human isolates outside LPLs included 56 clade G(vi) isolates and 19 non-G(vi) isolates. Based on the MCC tree from the primary analysis, none of the non-G(vi) human isolates were likely to have been closely related to an isolate from the Alberta cattle population, suggesting that all 19 were imported. As a proportion of all non-LPL human isolates, these 19 constituted 25.3%. While it may be possible that all clade G(vi) isolates were part of a local evolving lineage, it is also possible that exchange of both cattle and food from other locations was causing the regular importation of clade G(vi) strains and infections. Thus, we used the proportion of non-LPL human isolates outside the G(vi) clade to estimate the proportion of non-LPL human isolates within the G(vi) clade that were imported; i.e., 56 X 25.3% = 14. We then conducted a similar exercise for cattle isolates.

To contextualize our results in terms of ongoing human disease burden, we created a timed phylogeny using a constant, unstructured coalescent model of the 199 Alberta isolates from the primary analysis and the additional Alberta Health isolates (Figure 1). The two sets of sequences were combined and down-sampled, leaving 272 human and 84 cattle isolates (Supplemental Table S1). We identified LPLs as above, and leveraged the near-complete sequencing of isolates from 2018 and 2019 to calculate the proportion of reported human cases associated with LPLs.

Finally, we created a timed phylogeny of Alberta, U.S. and global from 1996 to 2019 to test whether the LPLs were linked to ancestors from locations outside Canada (Supplemental Table S1). Due to the size of this tree, we created both unstructured and structured versions. Clade A isolates were excluded due to their small number in Alberta and high level of divergence from other *E. coli* O157:H7 clades. Down-sampling was conducted separately by species and location. The phylogeny included 358 Alberta, 350 U.S., and 61 global isolates after down-sampling.

All BEAST2 analyses were run for 100,000,000 Markov chain Monte Carlo iterations or until all parameters converged with effective sample sizes >200, whichever was longer. Exact binomial 95% confidence intervals (CIs) were computed for proportions.

## Results

### Description of Isolates

Across the 1,215 isolates included in the analyses, we identified 12,273 core genome SNPs. Clade G(vi) constituted 73.6% (n=894) of all isolates (Figure 2a). Clade A, which is the most distinct of the *E. coli* O157:H7 clades, included non-Alberta isolates, 2 human isolates from Alberta, and no Alberta cattle isolates. The majority of all Alberta isolates belonged to the G(vi) clade (582 of 659; 88.3%), compared to 281 of the 1,560 (18.0%) randomly sampled U.S. PulseNet isolates that were successfully assembled and QCed. Among the 62 non-randomly sampled global isolates, only 2 (3.2%) were clade G(vi) (Figure 2b). There were 682 (76.3%) clade G(vi) isolates with the *stx1a/stx2a* profile and 210 (23.5%) with the *stx2a*-only profile, compared to 2 (0.6%) and 58 (18.1%), respectively, among the 321 isolates outside the G(vi) clade (Table 1).

**Figure 2.**
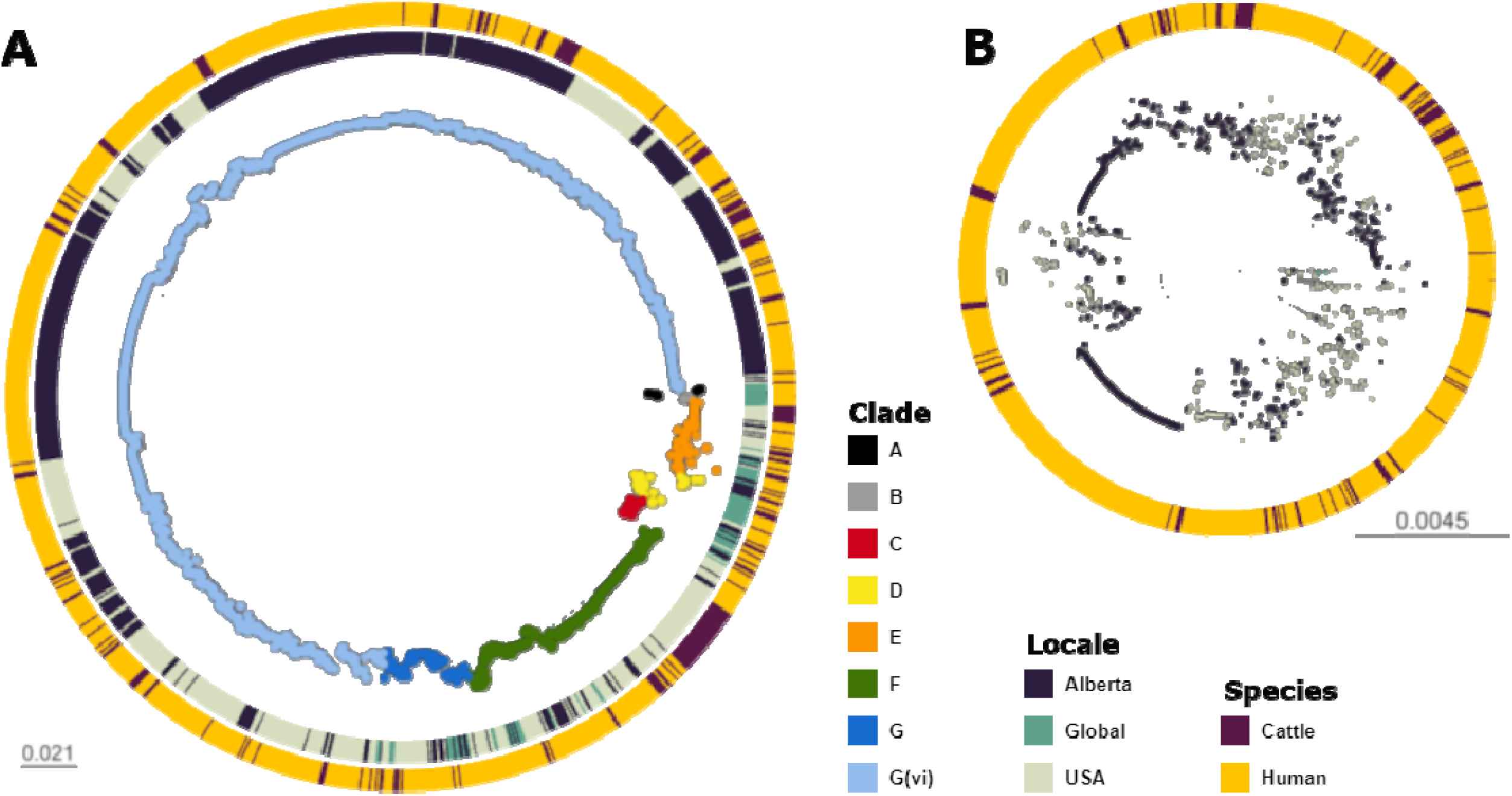
Maximum likelihood core SNP tree of the 1,215 *E. coli* O157:H7 isolates referenced in the study. This includes 659 isolates from Alberta, Canada, from 2007 through 2019, 494 isolates from the U.S. from 1996 through 2019, and 62 isolates from elsewhere around the globe from 2007 to 2016. The tree was rooted at clade A. (**A**) shows all clades, with tips colored according to clade, geographic origin shown on the inner ring, and species of origin on the outer ring. (**B**) shows clade G(vi), which constituted 73.6% of all isolates and 88.3% of Alberta isolates. Tips are colored by geographic origin, and the ring indicates species of origin.

**Table 1.**
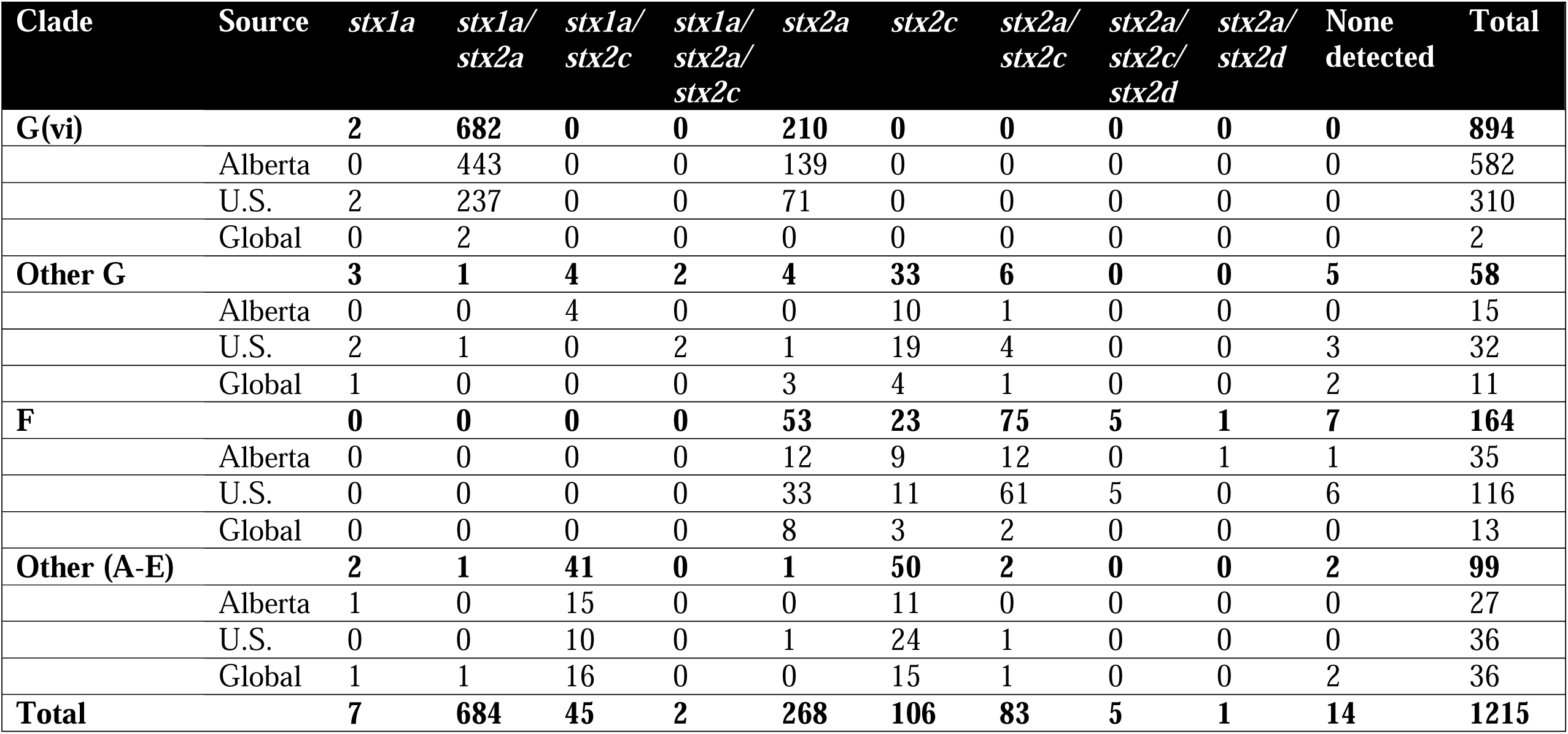
Distribution of study isolates by geographic source, clade, and Shiga toxin gene (*stx*) profile.

### The Majority of Clinical Cases Evolved from Local Cattle Lineages

In our primary sample of 121 human and 108 cattle isolates from Alberta from 2007 to 2015, SNP distances were comparable between species (Figure 3). Among sampled human cases, 19 (15.7%; 95% CI 9.7%, 23.4%) were within 5 SNPs of a sampled cattle strain. The median SNP distance between cattle sequences was 45 (IQR 36-56), compared to 54 (IQR 43-229) SNPs between human sequences from cases in Alberta during the same years.

**Figure 3.**
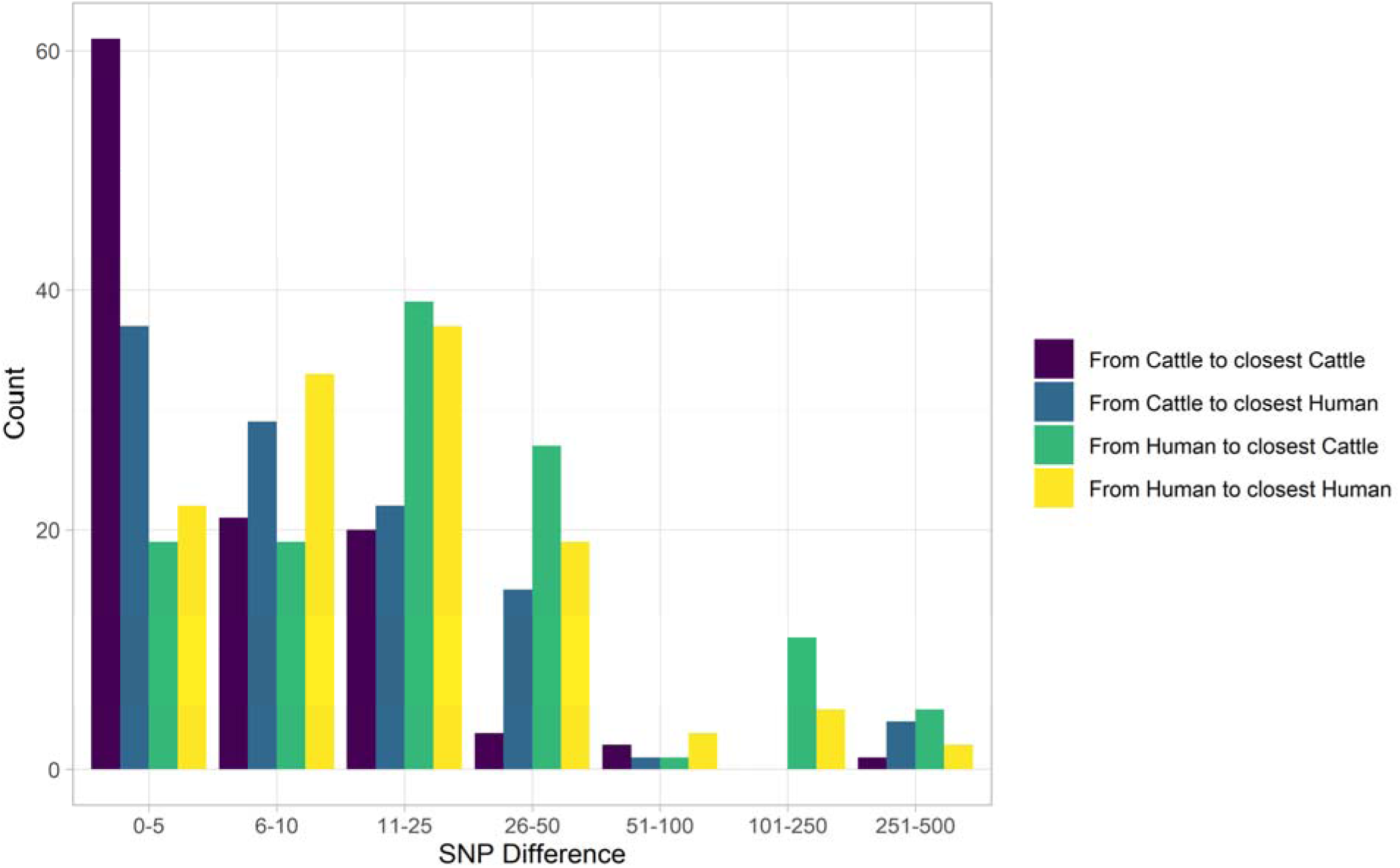
SNP distance between *E. coli* O157:H7 strains and their nearest relative, by species, Alberta, Canada, 2007-2015. Distances for isolates from 121 reported human cases and 108 beef cattle are shown. Cattle isolates were highly related with 56.5% of cattle isolates within 5 SNPs of another cattle isolate and 94.4% within 25 SNPs. Human isolates showed a bimodal distribution in their relationship to cattle isolates, with 86.0% within 50 SNPs of a cattle isolate and the remainder 185-396 SNPs apart. Nineteen human isolates (15.7%) were within 5 SNPs of a cattle isolate.

The phylogeny generated by our primary structured coalescent analysis indicated cattle were the primary reservoir, with a high probability that the hosts at nodes along the backbone of the tree were cattle (Figure 4). The root was estimated at 1802 (95% HPD 1731, 1861). The most recent common ancestor (MRCA) of clade G(vi) strains in Alberta was inferred to be a cattle strain, dated to 1969 (95% HPD 1959, 1979). With our assumption of a relaxed molecular clock, the mean clock rate for the core genome was estimated at 9.65x10^-5^ (95% HPD 8.13x10^-5^, 1.13x10^-4^) substitutions/site/year. The effective population size, *N_e_*, of the human *E. coli* O157:H7 population was estimated as 1060 (95% HPD 698, 1477), and for cattle as 73 (95% HPD 50, 98).

**Figure 4.**
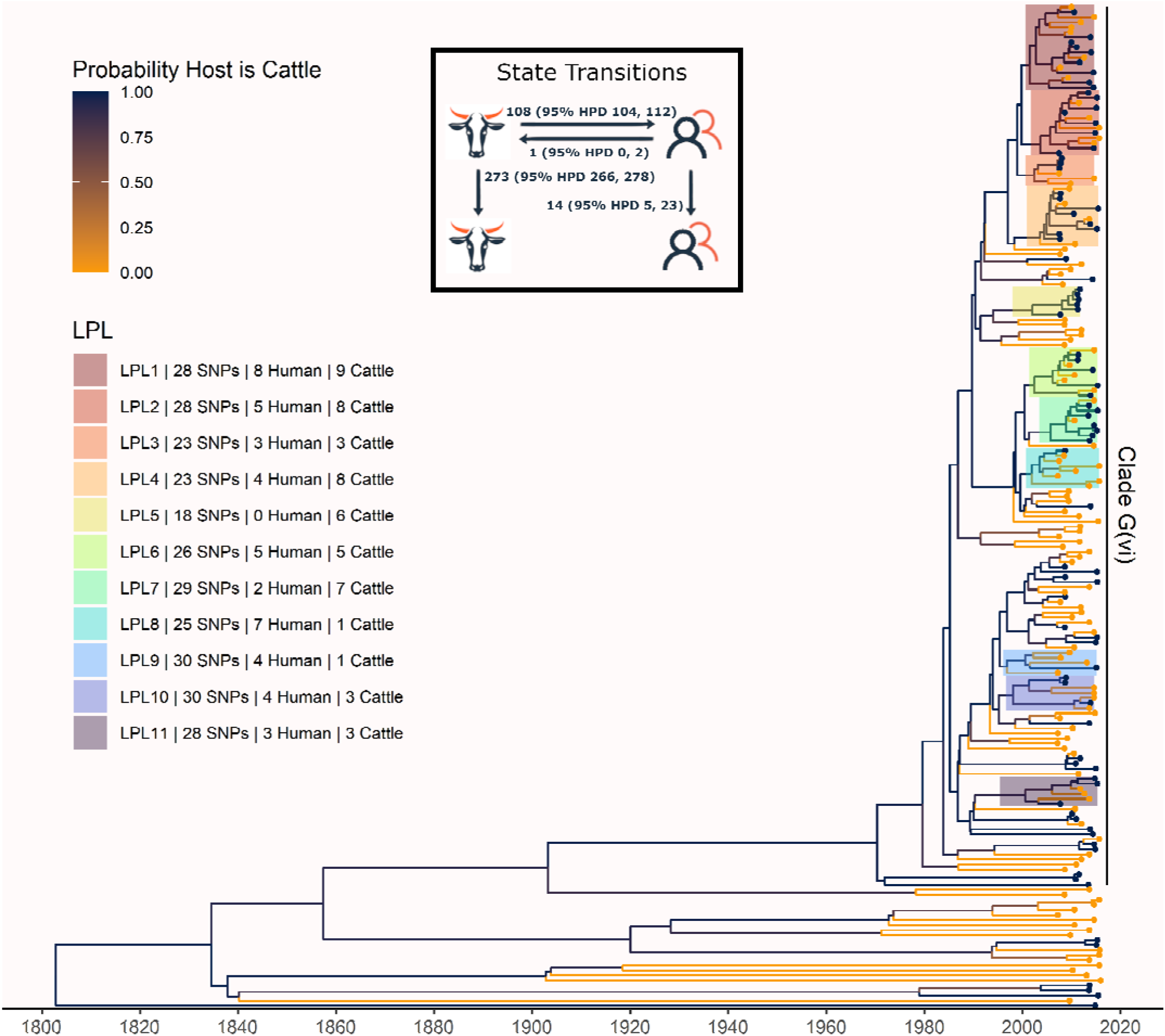
Maximum clade credibility (MCC) tree of structured coalescent analysis of *E. coli* O157:H7 strains isolated from 115 reported human cases and 84 beef cattle in Alberta, Canada, 2007-2015. Isolates were down-sampled prior to phylodynamic analysis to remove isolates that were highly similar. The structured coalescent analysis estimated migration and state transitions between humans and cattle. The MCC tree was colored by inferred host, cattle (blue) or human (orange). The minimum SNP distance between all isolates in the LPL, and the number of human and cattle isolates after down-sampling are shown for each LPL. The majority of ancestral nodes inferred as cattle suggests cattle as the primary reservoir. The root was estimated at 1802 (95% HPD 1731, 1861). Eleven locally persistent lineages (LPLs) were identified, all in the G(vi) clade and labeled LPL 1 through 11. For each LPL, the minimum number of SNPs all isolates are within (30 was set as the maximum) and the number of human and cattle isolates within the LPL, not including down-sampled isolates, are shown. With down-sampled isolates reincorporated, LPLs accounted for 46 human (38.0%) and 71 cattle (65.7%) isolates. The structured coalescent model estimated 108 cattle-to-human state transitions between branches, compared to only 14 human-to-human transitions, inferring cattle as the origin of 88.5% of human lineages.

We estimated 108 (95% HPD 104, 112) human lineages arose from cattle lineages, and 14 (95% HPD 5, 23) arose from other human lineages (Figure 4). In other words, 88.5% of human lineages seen in Alberta from 2007 to 2015 arose from cattle lineages. We observed minimal influence of our choice of priors (Supplemental Figure S2). Our sensitivity analysis of equal numbers of isolates from cattle and humans was largely consistent with our primary results, estimating that 94.3% of human lineages arose from cattle lineages (Supplemental Figure S3).

### Locally Persistent Lineages Account for the Majority of Ongoing Human Disease

In our primary analysis, we identified 11 locally persistent lineages (LPLs) (Figure 4). After reincorporating down-sampled isolates, LPLs included a range of 5 (G(vi)-AB LPL 9) to 26 isolates (G(vi)-AB LPL 1), with an average of 10. LPL assignment was based on the MCC tree of the combination of four independent chains. LPLs persisted for 5-9 years, with the average LPL spanning 8 years. By definition, MRCAs of each LPL were required to have a posterior probability ≥95% on the MCC tree, and in practice all had posterior probabilities of 99.7-100%. Additionally, examining all trees sampled from the four chains supported the same major lineages (Supplemental Figure S4). Our sensitivity analysis of equal numbers of isolates from cattle and humans identified 10 of the same 11 LPLs (Supplemental Figure S3). G(vi)-AB LPL 9 was no longer identified as an LPL, because it fell below the 5-isolate threshold after subsampling. Additionally, G(vi)-AB LPL 8 expanded to include a neighboring branch.

LPLs tended to be clustered on the MCC tree. G(vi)-AB LPLs 1-4, 6-8, and 9 and 10 were clustered with MRCAs inferred at 1996 (95% HPD 1992, 1999), 1998 (95% HPD 1995, 2000), and 1993 (95% HPD 1989, 1996), respectively (Figure 4). Cattle were the inferred host of all three ancestral nodes. LPLs were assigned using a threshold of 30 SNPs. In sensitivity analysis testing alternate SNP thresholds, we observed LPLs mimicking the larger clusters of the LPLs from our primary analysis (Supplemental Figure S5).

LPLs included 71 of 108 (65.7%; 95% CI 56.0%, 74.6%) cattle and 46 of 121 (38.0%; 95% CI 29.3%, 47.3%) human isolates. Of the remaining human isolates, 33 (27.3%) were associated with imported infections and 42 (34.7%) with infections from transient local strains. Of the remaining cattle isolates, 11 (10.2%) were imported and 26 (24.1%) were associated with transmission from transient strains. Of the 117 isolates in LPLs, 7 (6.0%) carried only *stx2a*, and the rest *stx1a/stx2a*. Among the 112 non-LPL isolates, 1 (0.9%) was *stx1a*-only, 27 (24.1%) were *stx2a*-only, 5 (4.5%) were *stx2c*-only, 68 (60.7%) were *stx1a/stx2a*, 6 (5.4%) were *stx1a/stx2c*, and 5 (4.5%) were *stx2a/stx2c*.

To understand long-term persistence, we expanded the phylogeny with additional Alberta Health isolates from 2009 to 2019 (Supplemental Table S1). Six of the 11 LPLs identified in our primary analysis, G(vi)-AB LPLs 1, 2, 4, 7, 10, and 11, continued to cause disease during the 2016 to 2019 period (Figure 5). With most cases reported during 2018 and 2019 sequenced, we were able to estimate the proportion of reported *E. coli* O157:H7 associated with LPLs. Of 217 sequenced cases reported during these two years, 162 (74.7%; 95% CI 68.3%, 80.3%) arose from Alberta LPLs. The *stx* profile of LPL isolates shifted as compared to the primary analysis, with 83 (51.2%; 95% CI 43.3%, 59.2%) of the LPL isolates encoding only *stx2a* and the rest *stx1a/stx2a* (Figure 6). Among the 55 non-LPL isolates during 2018-2019, the *stx2c*-only profile emerged with 16 (29.1%; 95% CI 17.6%, 42.9%) isolates, *stx2a*-only was found in 6 (10.9%; 95% CI 4.1%, 22.2%) isolates, and 5 (9.1%; 95% CI 3.0%, 20.0%) isolates carried both *stx2a* and *stx2c*.

**Figure 5.**
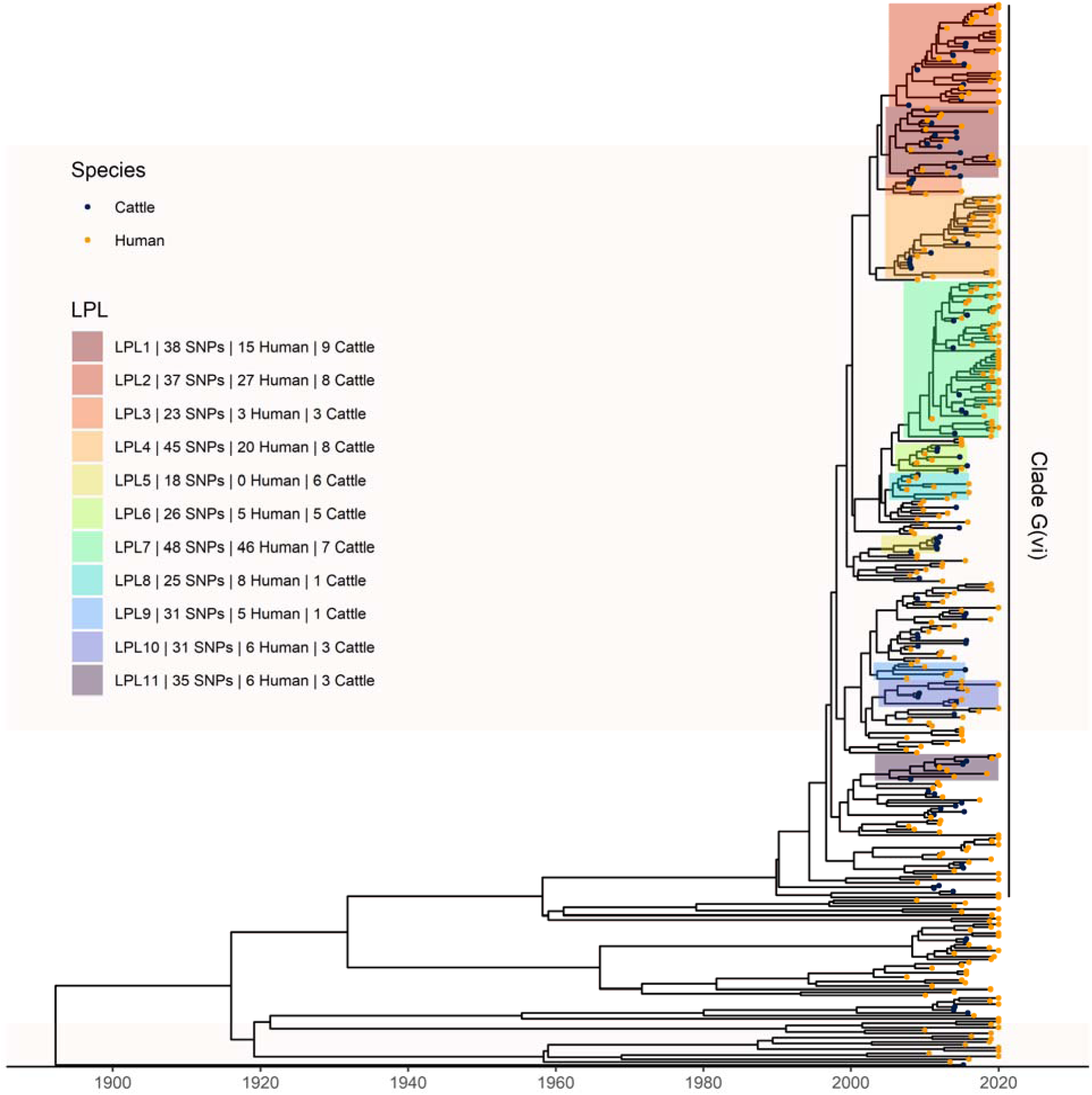
Extension of Alberta, Canada *E. coli* O157:H7 analysis to include 229 randomly selected study isolates and 430 additional public health isolates available from 2009 to 2019. The maximum clade credibility (MCC) tree was constructed from a coalescent analysis with constant population size after down-sampling. Six locally persistent lineages (LPLs) in clade G(vi) continued to be associated with disease after the initial study period. LPLs are colored and labeled as in Figure 4. After re-incorporating the down-sampled isolates, 74.7% of reported cases in 2018 and 2019 were associated with an LPL.

**Figure 6.**
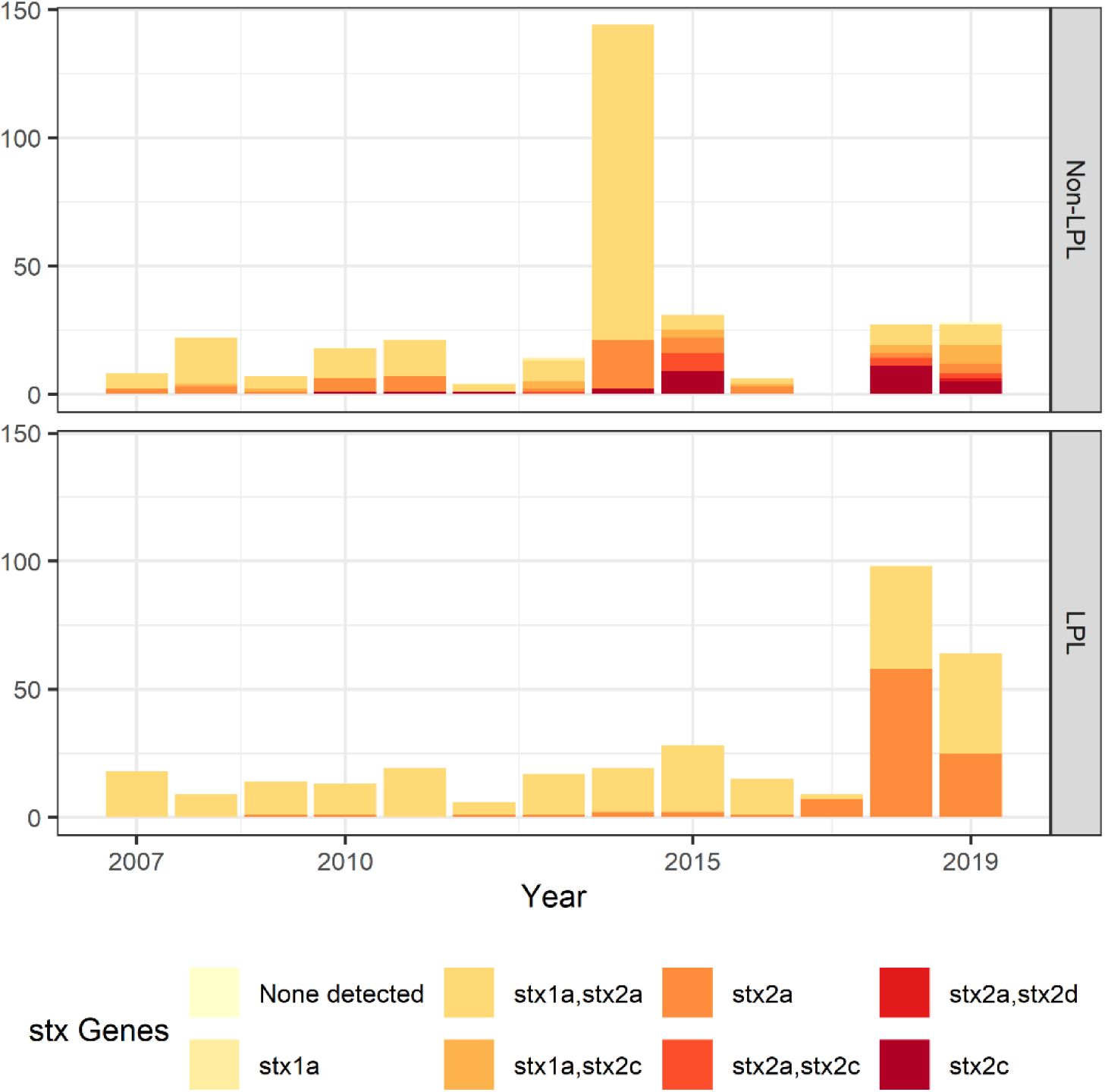
Shiga toxin gene (*stx*) profile by locally persistent lineage (LPL) status of extended analysis of Alberta, Canada *E. coli* O157:H7 isolated from cattle and humans, 2007 to 2019. The *stx* profile across all clades shifted from the initial study period (2007-2015) to the later study period (2016-2019), with more of the virulent *stx2a*-only profile observed in 2018 and 2019 than in previous years. In 2018 and 2019, 51.2% of LPL isolates carried only *stx2a*, compared to 10.9% of non-LPL isolates. The peak in sequences in 2014 is due to two outbreaks; routine sequencing began in 2018 and 2019, accounting for the rise in sequenced cases during those years.

All 5 large (≥10 cases) sequenced outbreaks in Alberta during the study period were within clade G(vi). G(vi)-AB LPLs 2 and 7 gave rise to 3 large outbreaks, accounting for 117 cases (both sequenced and unsequenced), including 83 from an extended outbreak by a single strain in 2018 and 2019, defined as isolates within 5 SNPs of one another. The two large outbreaks that did not arise from LPLs both occurred in 2014 and were responsible for 164 cases.

### Locally Persistent Lineages Were Not Imported

Of the 494 U.S. isolates analyzed, 9 (1.8%; 95% CI 0.8%, 3.4%) occurred within Alberta LPLs after re-incorporating down-sampled isolates (Figure 7). None of the 62 global isolates were associated with Alberta LPLs. The 9 U.S. isolates were part of G(vi)-AB LPLs 2 (n=3), 4 (n=4), 7 (n=1), and 11 (n=1), all of which had Alberta isolates that spanned 9 to 13 years and predated the U.S. isolates. There was no evidence of U.S. or global ancestors of LPLs. Based on migration events calculated from the structured tree, we estimated that 11.0% of combined human and cattle Alberta lineages were imported (Supplemental Table S2). Alberta sequences were separated from U.S. and global sequences by a median of 63 (IQR 45-236) and 225 (IQR 209-249) SNPs, respectively.

**Figure 7.**
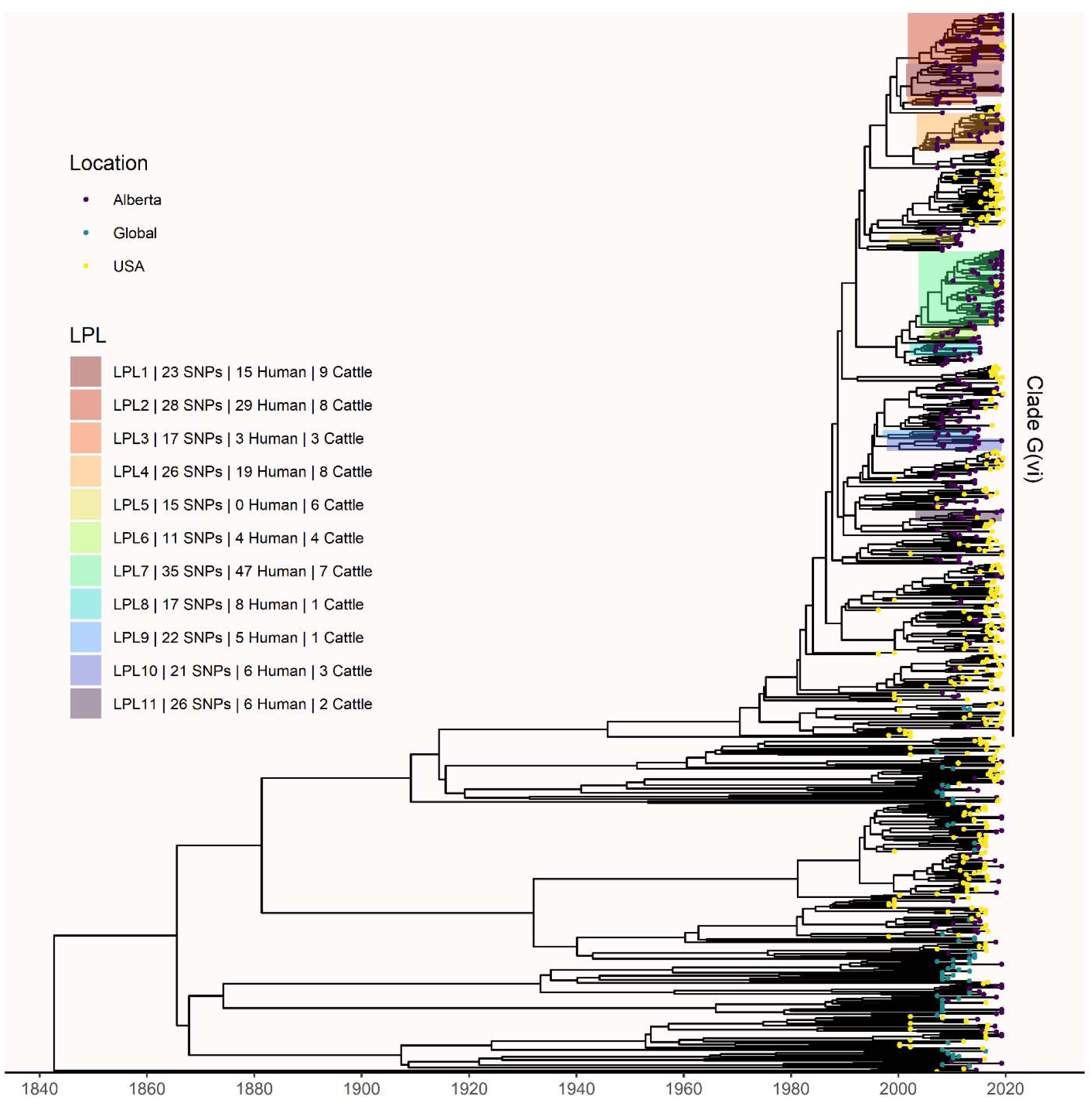
Comparison of Alberta, U.S., and global *E. coli* O157:H7 isolates. Tips are colored based on isolate origin, and LPLs from Figure 4 are highlighted. A total of nine U.S. isolates arose from Alberta LPLs 2, 4, 7, and 11, all of which had Alberta isolates predating the U.S. isolates. No global isolates were associated with Alberta LPLs. Clade A was excluded from the analysis due to its high level of divergence from the rest of the *E. coli* O157:H7 population.

Including U.S. and global isolates in the phylogeny did not change which LPLs we identified (Figure 7). The minimum SNPs that LPL isolates differed by was lower than in the Alberta-only analyses, because the core genome shared by all Alberta, U.S., and global isolates was smaller than that of only the Alberta isolates. Alberta sequences included in some LPLs changed slightly. G(vi)-AB LPL 4 lost 3 Alberta isolates from clinical cases, and G(vi)-AB LPLs 6 and 11 both lost 1 cattle and 1 human isolate from Alberta. In these LPLs, the isolates no longer included were the most outlying isolates in the LPLs defined using only Alberta isolates (Figure 5). Of the 217 Alberta human isolates from 2018 and 2019, 160 (73.7%) were still associated with LPLs after the addition of U.S. and global isolates, demonstrating stability of the extended analysis results.

## Discussion

Focusing on a region that experiences an especially high incidence of STEC, we conducted a deep genomic epidemiologic analysis of *E. coli* O157:H7’s multi-host disease dynamics. Our study identified multiple locally evolving lineages transmitted between cattle and humans. These were persistently associated with *E. coli* O157:H7 illnesses over periods of up to 13 years, the length of our study. Of clinical importance, there was a dramatic shift in the *stx* profile of the strains arising from locally persistent lineages toward strains carrying only *stx2a*, which has been associated with increased progression to hemolytic uremic syndrome (HUS).^27^

Our study has provided quantitative estimates of cattle-to-human migration in a high incidence region, the first such estimates of which we are aware. Our estimates are consistent with prior work that established an increased risk of STEC associated with living near cattle.^2,12–16^ We showed that 88.5% of strains infecting humans arose from cattle lineages. These transitions can be seen as a combination of the infection of humans from local cattle or cattle-related reservoirs in clade G(vi) and the historic evolution of *E. coli* O157:H7 from cattle in the rare clades. While our findings indicate the majority of human cases arose from cattle lineages, transmission may involve intermediate hosts or environmental sources several steps removed from the cattle reservoir. Small ruminants (e.g., sheep, goats) have been identified as important STEC reservoirs,^13,16,26^ and Alberta has experienced outbreaks linked to swine.^8^ Exchange of strains between cattle and other animals may occur if co-located, if surface water sources near farms are contaminated, and through wildlife, including deer, birds, and flies.^28,29^ Humans can also become infected from environmental sources, such as through swimming in contaminated water.

Although transmission systems may be multi-faceted, our analysis demonstrates that local cattle remain an integral part of the transmission system for the vast majority of cases, even when they may not be the immediate source of infection. Indeed, despite our small sample of *E. coli* O157:H7 isolates from cattle, 15.7% of our human cases were within 5 SNPs of a cattle isolate, suggesting that cattle were a recent source of transmission, either through direct contact with the animal or their environments or consumption of contaminated food products.

The cattle-human transitions we estimated were based on structured coalescent theory, which we used throughout our analyses. This approach is similar to other phylogeographic methods that have previously been applied to *E. coli* O157:H7.^21^ We inferred the full backbone of the Alberta *E. coli* O157:H7 phylogeny as arising from cattle, consistent with the postulated global spread of the pathogen via ruminants.^21^ Our estimate of the origin of the serotype, at 1802 (95% HPD 1731, 1861), was somewhat earlier than previous estimates, but consistent with global (1890; 95% HPD 1845, 1925)^21^ and United Kingdom (1840; 95% HPD 1817, 1855)^30^ studies that used comparable methods. Our dating of the G(vi) clade in Alberta to 1969 (95% HPD 1959, 1979) also corresponds to proposed migrations of clade G into Canada from the U.S. in 1965-1977.^21^ Our study thus adds to the growing body of work on the larger history of *E. coli* O157:H7, providing an in-depth examination of the G(vi) clade.

Our identification of the 11 locally persistent lineages (LPLs) is significant in demonstrating that the majority of Alberta’s reported *E. coli* O157:H7 illnesses are of local origin. Our definition ensured that every LPL had an Alberta cattle strain and at least 5 isolates separated by at least 1 year, making the importation of the isolates in a lineage highly unlikely. For an LPL to be fully imported, cattle and human isolates would need to be repeatedly imported from a non-Alberta reservoir where the lineage was persisting over several years. Further supporting the evolution of the LPLs within Alberta, all 11 LPLs were in clade G(vi), several were phylogenetically related with MRCAs dating to the late 1990s, and few non-Alberta isolates fell within LPLs. The 9 U.S. isolates associated with Alberta LPLs may reflect Alberta cattle that were slaughtered in the U.S. or infections in travelers from the U.S. Thus, we are confident that the identified LPLs represent locally evolving lineages and potential persistent sources of disease. We also showed that the identification of these LPLs was robust to the sampling strategy, with only the smallest LPL failing to be identified after subsampling left it with <5 isolates.

We estimated the proportion of *E. coli* O157:H7 that were imported into Alberta in two ways. Based on our LPL analysis, we estimated only 27% of human and 10% of cattle *E. coli* O157:H7 isolates were imported. This was slightly higher than the overall importation estimate of 11% for all Alberta lineages from our global structured coalescent analysis. Our global structured coalescent analysis also estimated that 3% of lineages in the U.S. and 2% of lineages outside the U.S. and Canada had been exported from Alberta, suggesting that Alberta is not a significant contributor to global *E. coli* O157:H7 burden beyond its borders. These results place the *E. coli* O157:H7 population in Alberta within a larger context, indicating that the majority of disease can be considered local. At least one study has attempted to differentiate local vs. non-local lineages based on travel status,^31^ which may be appropriate in some locations but can miss cases imported through food products, such as produce imported from other countries. To our knowledge, our study provides the first comprehensive determination of local vs. imported status for *E. coli* O157:H7 cases using external reference cases. Similar studies in regions of both high and moderate incidence would provide further insight into the role of localization on *E. coli* O157:H7 incidence.

Of the 11 lineages we identified as LPLs during the 2007-2015 period, 6 were also associated with cases that occurred during the 2016-2019 period. During the initial period, 38% of human cases were linked to an LPL, and 6% carried only *stx2a*. The risk of HUS increases in strains of STEC carrying only *stx2a*, relative to *stx1a/stx2a*,^27^ meaning the earlier LPL population had fewer high-virulence strains. In 2018 and 2019, the 6 long-term LPLs were associated with both greater incidence and greater virulence, encompassing 75% of human cases with more than half of LPL isolates carrying only *stx2a*. The cause of this shift remains unclear, though shifts toward greater virulence in *E. coli* O157:H7 populations have been seen elsewhere.^32^ The growth and diversity of G(vi)-AB LPLs 2, 4, and 7 in the later period suggest these lineages were in stable reservoirs or adapted easily to new niches. Identifying these reservoirs could yield substantial insights into the disease ecology that supports LPL maintenance and opportunities for disease prevention, given the significant portion of illnesses caused by persistent strains.

The high proportion of cases associated with cattle-linked local lineages is consistent with what is known about the role of cattle in STEC transmission. Among sporadic STEC infections, 26% have been attributed to animal contact and the farm environment, with a further 19% to pink or raw meat.^11^ Similarly, 24% of *E. coli* O157 outbreaks in the U.S. have been attributed to beef, animal contact, water, or other environmental sources.^10^ In Alberta, these are all inherently local exposures, given that 90% of beef consumed in Alberta is produced and/or processed there. Even person-to-person transmission, responsible for 15% of sporadic cases and 16% of outbreaks,^10,11^ includes secondary transmission from cases infected from local sources, which may explain our estimate of 11.5% of human lineages arising from other human lineages.

We developed a novel measure of persistence for use in this study, specifically for the purposes of identifying lineages that pose an ongoing threat to public health in a specific region.

Persistence has been variably defined in the literature, for example as shedding of the same strain for at least 4 months.^33^ Most recently, the U.S. CDC identified the first Recurring, Emergent, and Persistent (REP) STEC strain, REPEXH01, an *E. coli* O157:H7 strain detected since 2017 in over 600 cases. REPEXH01 strains are within 21 allele differences of one another (https://www.cdc.gov/ncezid/dfwed/outbreak-response/rep-strains/repexh01.html), and REP strains from similar enteric pathogens are defined based on allele differences of 13 to 104. Given that we used high resolution SNP analysis rather than cgMLST, we used a difference of ≤30 SNPs to define persistent lineages. While both our study and the REPEXH01 strain identified by CDC indicate that persistent strains of *E. coli* O157:H7 exist, the O157:H7 serotype was defined as sporadic in a German study using the 4-month shedding definition.^33^ This may be due to strain differences between the two locations, but it might also indicate that persistence occurs at the host community level, rather than the individual host level. Understanding microbial drivers of persistence is an active field of research, with early findings suggesting a correlation of STEC persistence to the accessory genome and traits such as biofilm formation and nutrient metabolism.^33,34^ Our approach to studying persistence was specifically designed for longitudinal sampling in high-incidence regions and may be useful for others attempting to identify sources that disproportionately contribute to disease burden. Although we used data from the reservoir species to help define the LPLs in this study, we are testing alternate approaches that rely on only routinely collected public health data.

We limited our analysis to *E. coli* O157:H7 despite the growing importance of non-O157 STEC, as historical multi-species collections of non-O157 isolates are lacking. As serogroups differ meaningfully in exposures,^35^ our results may not be generalizable beyond the O157 serogroup. However, cattle are still believed to be a primary reservoir for non-O157 STEC, and cattle density is associated with risk of several non-O157 serogroups.^14^ Person-to-person transmission remains a minor contributor to STEC burden. For all of these reasons, if we were to conduct this analysis in non-O157 STEC, we expect the majority of human lineages would arise from cattle lineages. Additionally, persistence within the cattle reservoir has been observed for a range of serogroups,^33^ suggesting that LPLs also likely exist among non-O157 STEC. Our findings may have implications beyond STEC, as well. Other zoonotic enteric pathogens such as *Salmonella* and *Campylobacter* can persist, and outbreaks are regularly linked to localized animal populations and produce-growing operations contaminated by animal feces. The U.S. CDC has also defined REP strains for these pathogens. LPLs could shed light on how and where persistent strains are proliferating, and thus where they can be controlled.

The identification of LPLs serves multiple purposes, because they suggest the existence of local reservoir communities that maintain specific strains for long periods. First, they further our understanding of the complex systems that allow STEC to persist. In this study, the LPLs we identified persisted for 5 to 13 years. The reservoir communities that enable persistence could involve other domestic and wild animals previously found to carry STEC.^13,16,26,28,29,36^ The feedlot environment also likely plays an important role in persistence, as water troughs and pen floors have been identified as important sources of STEC for cattle.^37^ Identifying LPLs is a first step in identifying these reservoir communities and determining what factors enable strains to persist, so as to identify them for targeted interventions. Second, identification of these LPLs in cattle could identify the specific local reservoirs of STEC. Similar to source tracing in response to outbreaks, LPLs provide an opportunity for cattle growers to identify cattle carrying the specific strains that are associated with a large share of human disease in Alberta. While routinely vaccinating against STEC has not been shown to be efficacious or cost effective,^38^ a ring-type vaccination strategy in response to an identified LPL isolate could overcome the limitations of previous vaccination strategies. Third, identification of new clinical cases infected with LPL strains could help direct public health investigations toward local sources of transmission. Finally, the disease burden associated with LPLs could be compared across locations and may help explain how high-incidence regions differ from regions with lower incidence.

Our analysis was limited to only cattle and humans. Had isolates from a wider range of potential reservoirs been available, we would have been able to elucidate more clearly the roles that various hosts and common sources of infection play in local transmission. Additional hosts may help explain the 1 human-to-cattle predicted transmission, which could be erroneous. As with all studies utilizing public health data, sampling from only severe cases of disease is biased towards clinical isolates. In theory, this could limit the genetic variation among human isolates if virulence is associated with specific lineages. However, clinical isolates were more variable than cattle isolates, dominating the most divergent clade A, so overrepresentation of severe cases does not appear to have appreciably biased the current study. Similarly, in initially selecting an equal number of human and cattle isolates, we sampled a larger proportion of the human-infecting *E. coli* O157:H7 population compared to the population that colonizes cattle. As cattle are the primary reservoir of *E. coli* O157:H7, the pathogen is more prevalent in cattle than in humans, who appear to play a limited role in sustained transmission. In sampling a larger proportion of the strains that infect humans, we likely sampled a wider diversity of these strains compared to those in cattle, which could have biased the analysis toward finding humans as the ancestral host. Thus, the proportion of human lineages arising from cattle lineages (88.5%) might be underestimated, which is also suggested by our sensitivity analysis of equal numbers of cattle and clinical isolates. Finally, we were not able to estimate the impact of strain migration between Alberta and the rest of Canada, because locational metadata for publicly available *E. coli* O157:H7 sequences from Canada was limited.

*E. coli* O157:H7 infections are a pressing public health problem in many high incidence regions around the world including Alberta, where a recent childcare outbreak caused >300 illnesses. In the majority of sporadic cases, and even many outbreaks,^10^ the source of infection is unknown, making it critical to understand the disease ecology of *E. coli* O157:H7 at a system level. Here we have identified a high proportion of human cases arising from cattle lineages and a low proportion of imported cases. Local transmission systems, including intermediate hosts and environmental reservoirs, need to be elucidated to develop management strategies that reduce the risk of STEC infection. In Alberta, local transmission is dominated by a single clade, and over the extended study period, persistent lineages caused an increasing proportion of disease. The local lineages with long-term persistence are of particular concern because of their increasing virulence, yet they also present opportunity as larger, more stable targets for reservoir identification and control.

## Acknowledgements

We would like to acknowledge Dr. Angela Ma, Hannah Tyrrell, and Dr. Surangi Thilakarathna for their work preparing clinical isolates for sequencing, and Dr. Jesse Berman for reviewing an early version of this manuscript. The authors would like to acknowledge and thank the PulseNet participating laboratories, whose data was used for the creation of this publication.

## Author Contributions

**Gillian A.M. Tarr**: Conceptualization, Methodology, Software, Formal analysis, Data curation, Writing - Original Draft, Visualization, Funding acquisition. **Linda Chui**: Conceptualization, Methodology, Resources, Writing - Review & Editing, Supervision, Funding acquisition. **Kim Stanford**: Conceptualization, Methodology, Resources, Writing - Review & Editing, Supervision, Funding acquisition. **Emmanuel W. Bumunang**: Investigation, Data curation, Writing - Review & Editing. **Rahat Zaheer**: Methodology, Investigation, Writing - Review & Editing. **Vincent Li**: Validation, Data curation, Writing - Review & Editing. **Stephen B. Freedman**: Conceptualization, Writing - Review & Editing, Project administration, Funding acquisition. **Chad Laing**: Conceptualization, Methodology, Writing - Review & Editing, Funding acquisition. **Tim A. McAllister**: Conceptualization, Methodology, Resources, Writing - Review & Editing, Supervision, Project administration, Funding acquisition.

## Data Sharing Statement

Data from this study not previously published will be made available at publication. Deidentified participant data, associated NCBI accession numbers for sequence data, and an accompanying data dictionary is provided as an attached data supplement.

## Supplemental Information

### STEC Case Definition

Alberta Health defines a confirmed case of Shiga toxin-producing *E. coli* (STEC), including *E. coli* O157:H7, as STEC isolation or Shiga toxin antigen or nucleic acid detection. Clinical illness, which may include diarrhea, bloody diarrhea, abdominal cramps, hemolytic uremic syndrome, thrombocytopenia purpura, or pulmonary edema, may or may not be present.

### Sampling from the Prior Distribution

Results in our Bayesian phylodynamic analyses are drawn from posterior distributions, which are influenced by both the data and the prior information we have about the system (Supplemental Table S1). In order to confirm that our primary results were not overly influenced by our prior assumptions, we conducted an analysis in which the sampling draws were made from the prior distribution, as opposed to the posterior distribution. We graphed these results against the sampling draws made from the posterior distributions from the four runs conducted for our primary analysis (each performed with a different random seed). The comparison shows that the draws from prior distribution differ markedly from the draws from the posterior distributions for the model’s key parameters (Supplemental Figure S2). From this, we concluded that our prior assumptions were not overly influencing the results of the primary analysis.

## Supplemental Figures

**Supplemental Figure S1.**
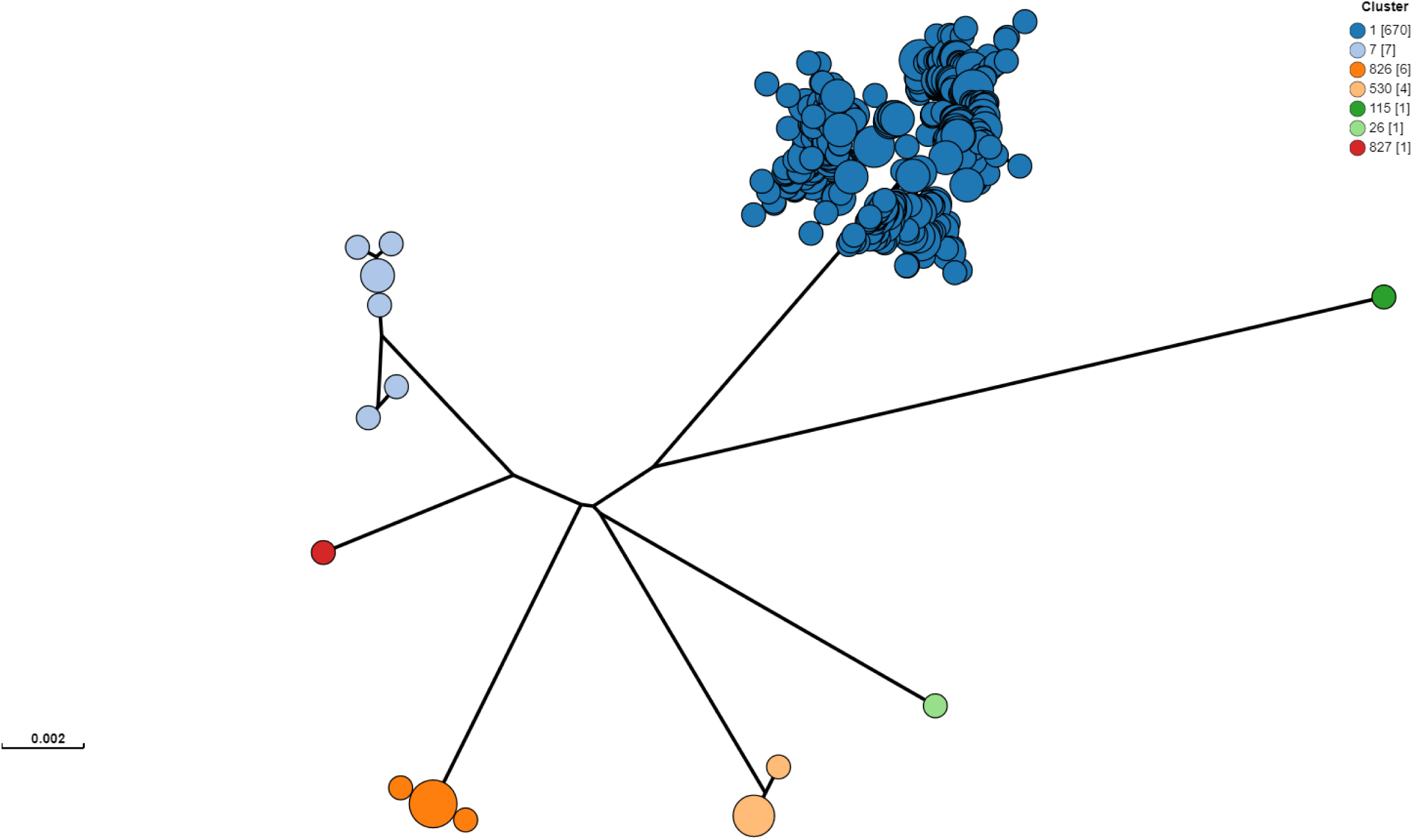
Genomic clustering of 690 Alberta *E. coli* O157 isolates. Clustering performed from raw reads using PopPUNK v2.5.0 with 10,146 *E. coli* reference genomes.^23^ From the 246 isolates selected for sequencing for the study and 445 additional Alberta Health isolates included for contextualization, one isolate was removed prior to clustering analysis, because it was identified through metadata review as an environmental (non-human, non-cattle) isolate. Cluster 1 included the Sakai and EDL933 reference strains. Clusters 826 and 827 were novel clusters. Isolates outside of Cluster 1 were excluded from all subsequent analyses.

**Supplemental Figure S2.**
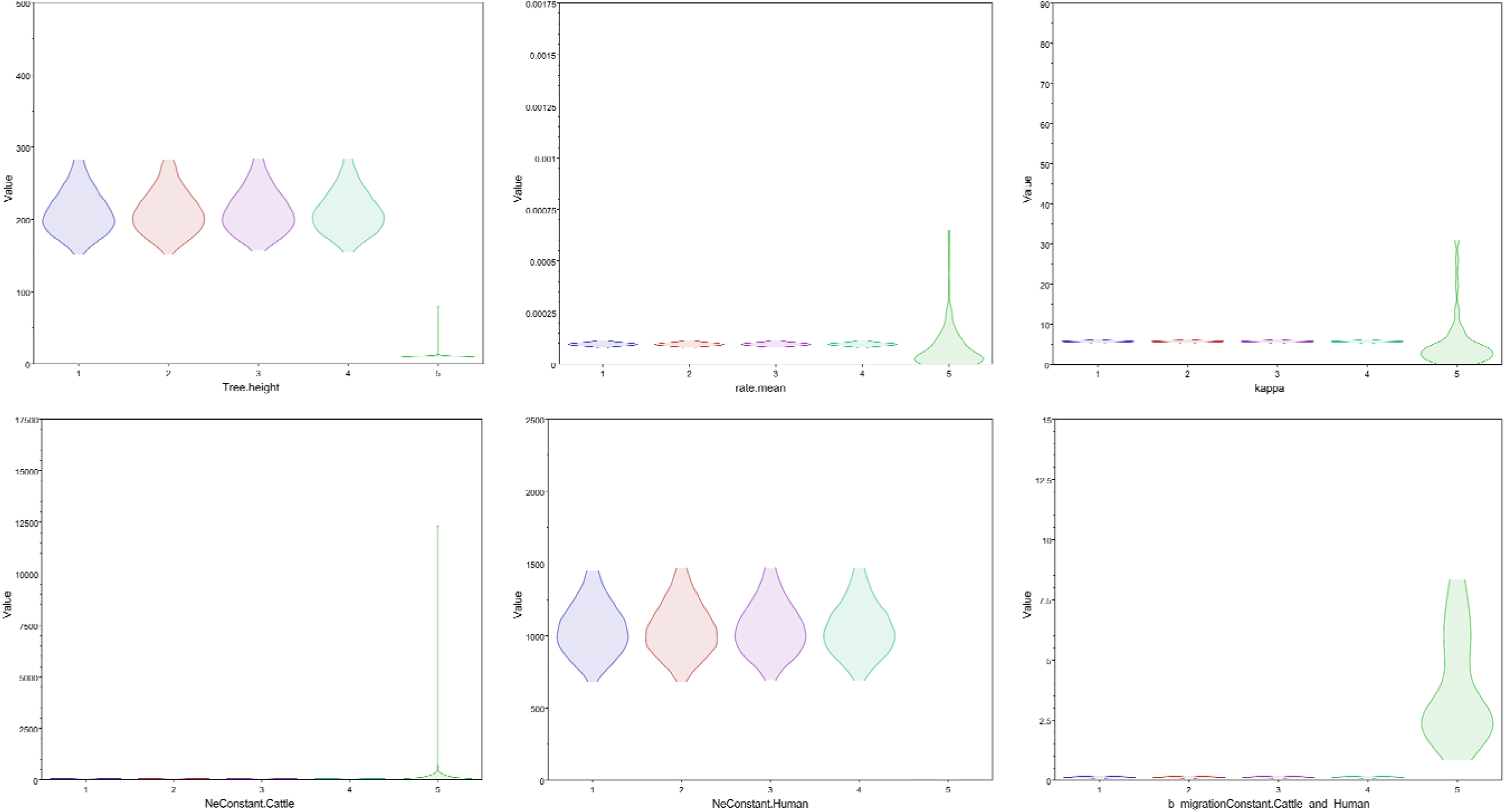
Key parameters drawn from the posterior distributions of four Markov chain Monte Carlo chains using different starting seeds compared to draws from the prior distribution. The posterior distributions of the four chains for the tree height, clock rate, kappa, cattle effective population size, human effective population size, and backward migration rate all differed substantially from the prior distribution, shown on the far right in green for each graph.

**Supplemental Figure S3.**
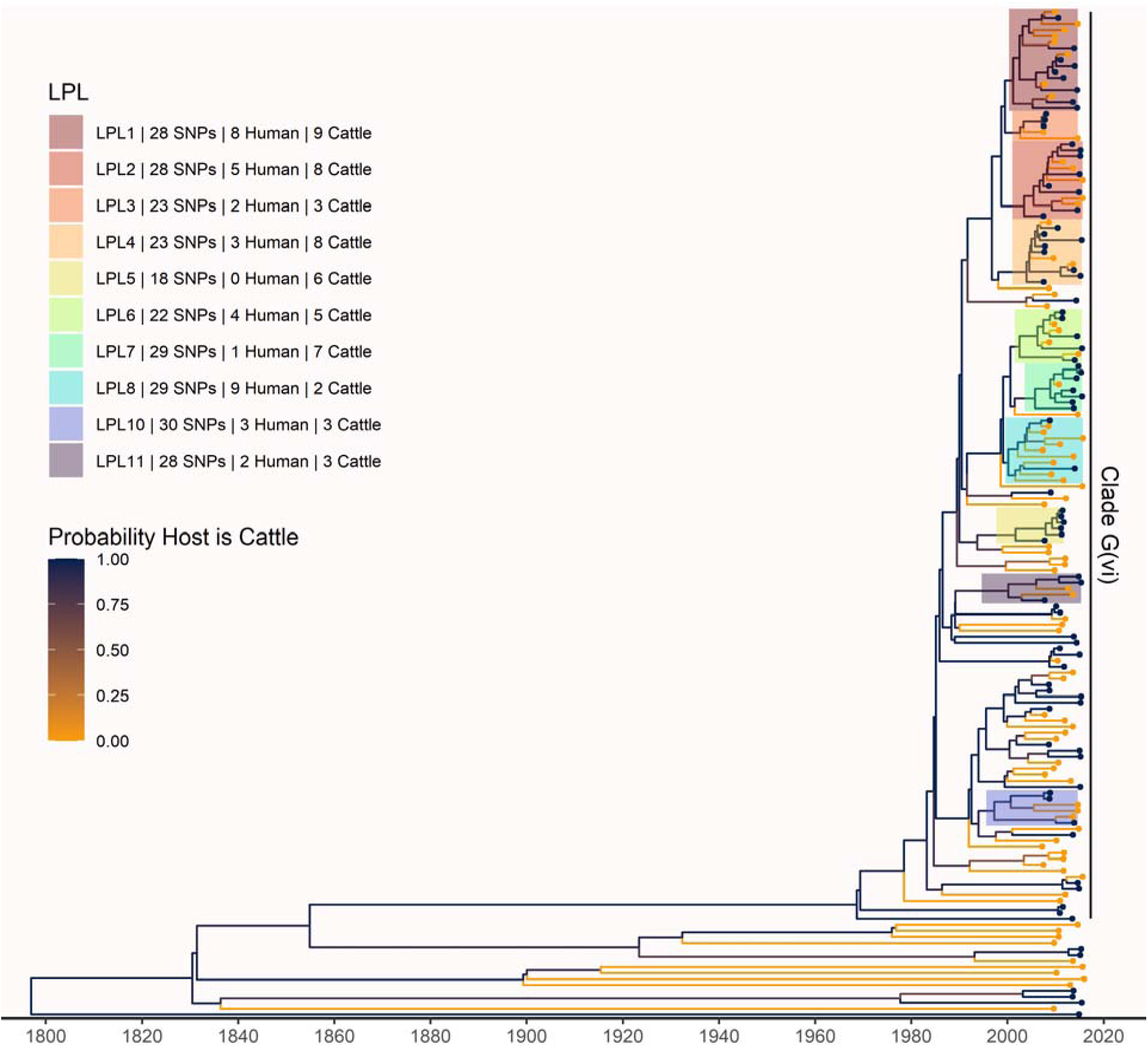
Maximum clade credibility (MCC) tree of structured coalescent analysis of 168 subsampled isolates from humans and cattle from Alberta, 2007-2015. LPLs are labeled as in the primary analysis (Figure 4). From an initial 121 human and 108 cattle isolates, our primary analysis contained 115 human and 84 cattle isolates after down-sampling. In this sensitivity analysis, we repeated our primary analysis with a subsample of 84 randomly selected isolates from humans and the 84 cattle isolates remaining after down-sampling. As in the primary analysis, cattle were inferred as the host of the majority of ancestral nodes. The root was estimated at 1796 (95% HPD 1722, 1859), very close to the root at 1802 estimated in the primary analysis. The most recent common ancestor (MRCA) of clade G(vi) strains in Alberta was inferred to be a cattle strain, dated to 1968 (95% HPD 1956, 1979), compared to 1969 in the primary analysis. We estimated 82 (95% HPD 79, 84) human lineages arose from cattle lineages, and 5 (95% HPD 0, 11) arose from other human lineages, meaning we inferred that 94.3% of human lineages arose from cattle lineages, compared to 88.5% in the primary analysis. The LPLs identified in this sensitivity analysis were mostly identical to those identified in the primary analysis. Differences in the sensitivity analysis were that G(vi)-AB LPL 8 expanded to a larger set of isolates; G(vi)-AB LPL 9, which included only 5 isolates in the primary analysis, was no longer identified as an LPL, as it no longer met the 5-isolate criterion; and there were minor topological changes.

**Supplemental Figure S4.**
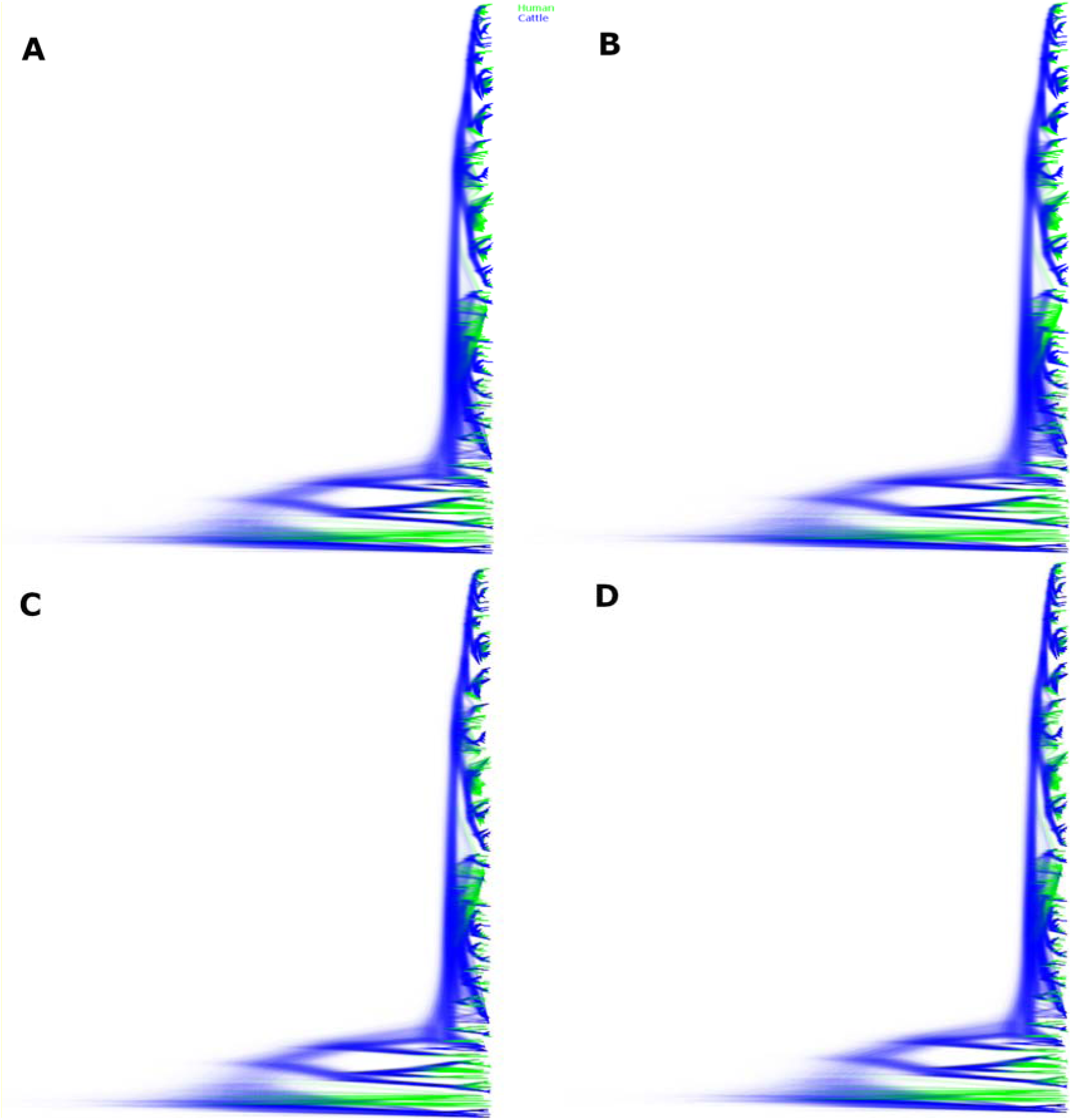
Sampled trees from four independent chains of the Alberta 2007-2015 structured coalescent analysis. Across the four chains, 1,070,460,000 trees were sampled. Depicted here are 963,414,000 post-burn-in trees, with panels A-D each showing the trees from one chain. Lineages inferred with cattle ancestry are shown in blue, lineages inferred with human ancestry are shown in green. Reflecting the strong support for the lineages identified as LPLs, the tree topology is well resolved.

**Supplemental Figure S5.**
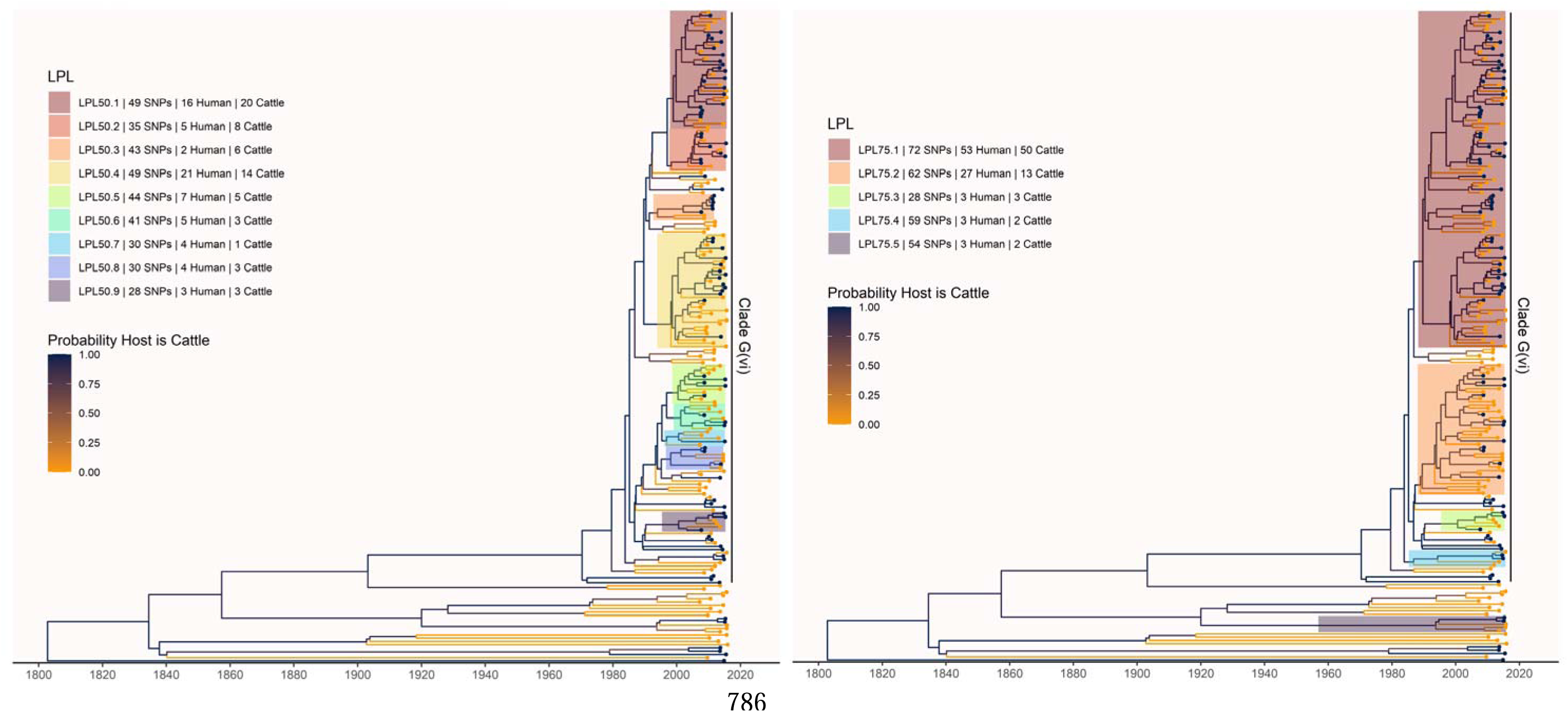
LPLs from the primary analysis defined using alternate SNP thresholds of 50 (left) and 75 (right) SNPs. LPL 50.1 includes G(vi)-AB LPLs 1, 2, and 3 from the primary analysis using a SNP threshold of 30. LPLs 50.2 and 50.3 correspond to G(vi)-AB LPLs 4 and 5, respectively. LPL 50.4 includes G(vi)-AB LPLs 6, 7, and 8. LPL 75.1 includes all of the above LPLs at the lower thresholds. All 50- and 75-threshold LPLs in this section of the tree include isolates not included at the lower SNP threshold(s). No part of LPLs 50.5 or 50.6 was identified as an LPL using a threshold of 30, but LPLs 50.7 and 50.8 correspond exactly to G(vi)-AB LPLs 9 and 10. LPL 75.2 includes LPLs 50.5-50.8 and G(vi)-AB LPLs 9 and 10. G(vi)-AB LPL 11, LPL 50.9, and LPL 75.3 are all identical. At the 75-SNP threshold, two additional LPLs are identified with no corresponding LPLs at the 30- or 50-SNP thresholds. One of these is outside the G(vi) clade.

## Supplemental Tables

**Supplemental Table S1.**
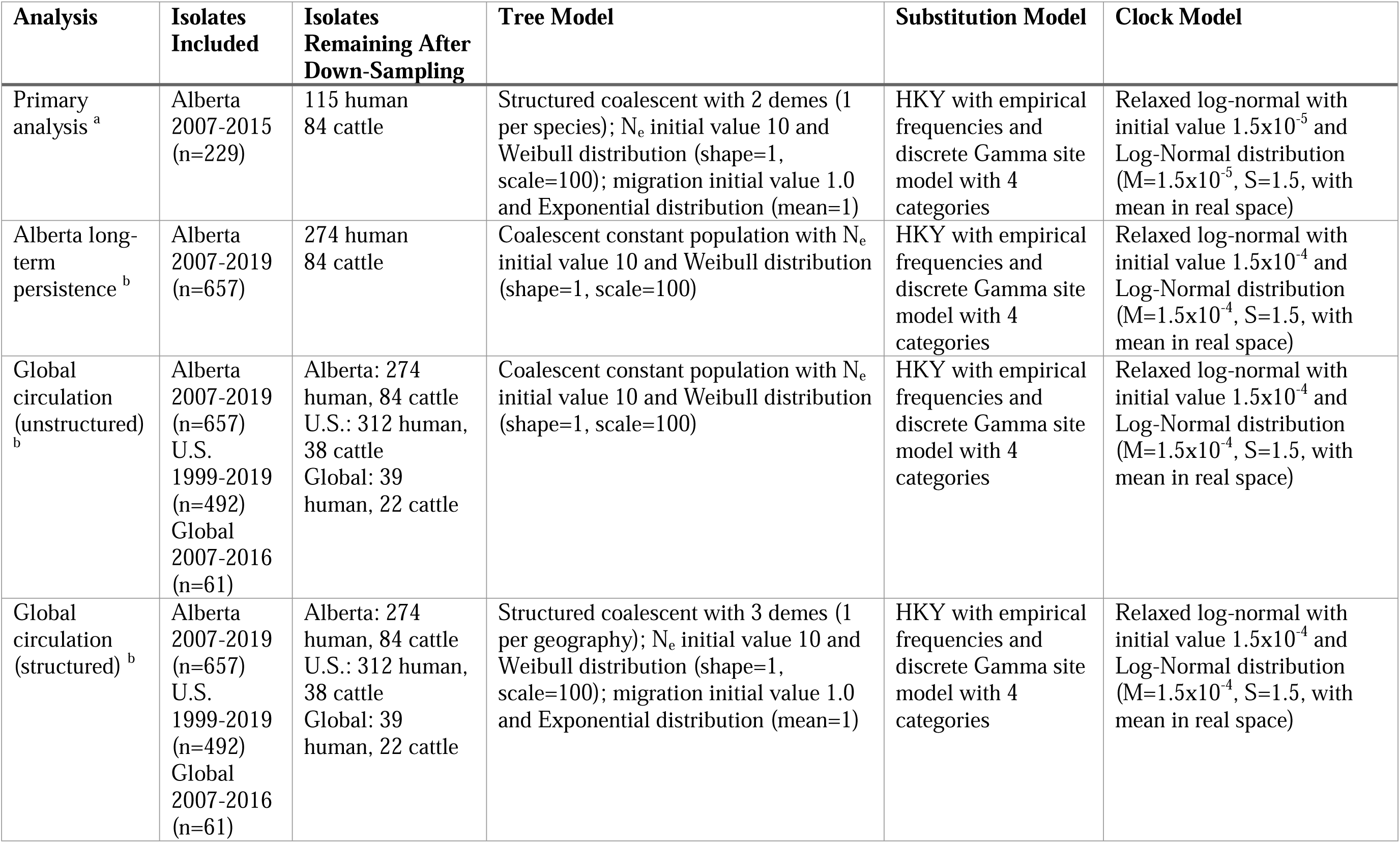
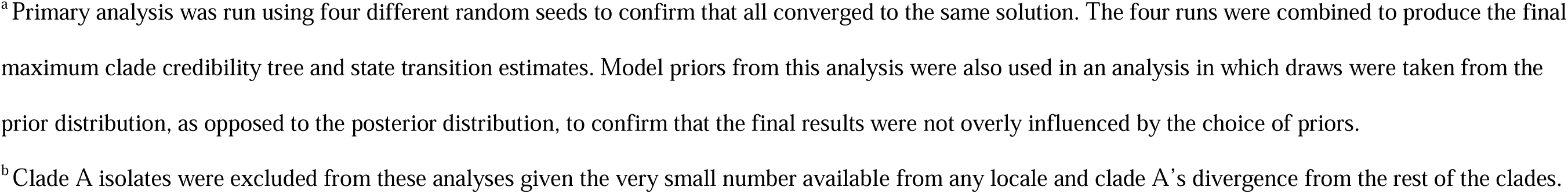
Analyses conducted and model priors.

| Analysis | Isolates Included | Isolates Remaining After Down-Sampling | Tree Model | Substitution Model | Clock Model |
| --- | --- | --- | --- | --- | --- |
| Primary analysis <sup>a</sup> | Alberta 2007-2015 (n=229) | 115 human<br>84 cattle | Structured coalescent with 2 demes (1 per species); $N_e$ initial value 10 and Weibull distribution (shape=1, scale=100); migration initial value 1.0 and Exponential distribution (mean=1) | HKY with empirical frequencies and discrete Gamma site model with 4 categories | Relaxed log-normal with initial value $1.5 \times 10^{-5}$ and Log-Normal distribution ($M=1.5 \times 10^{-5}$, $S=1.5$, with mean in real space) |
| Alberta long-term persistence <sup>b</sup> | Alberta 2007-2019 (n=657) | 274 human<br>84 cattle | Coalescent constant population with $N_e$ initial value 10 and Weibull distribution (shape=1, scale=100) | HKY with empirical frequencies and discrete Gamma site model with 4 categories | Relaxed log-normal with initial value $1.5 \times 10^{-4}$ and Log-Normal distribution ($M=1.5 \times 10^{-4}$, $S=1.5$, with mean in real space) |
| Global circulation (unstructured) <sup>b</sup> | Alberta 2007-2019 (n=657)<br>U.S. 1999-2019 (n=492)<br>Global 2007-2016 (n=61) | Alberta: 274 human, 84 cattle<br>U.S.: 312 human, 38 cattle<br>Global: 39 human, 22 cattle | Coalescent constant population with $N_e$ initial value 10 and Weibull distribution (shape=1, scale=100) | HKY with empirical frequencies and discrete Gamma site model with 4 categories | Relaxed log-normal with initial value $1.5 \times 10^{-4}$ and Log-Normal distribution ($M=1.5 \times 10^{-4}$, $S=1.5$, with mean in real space) |
| Global circulation (structured) <sup>b</sup> | Alberta 2007-2019 (n=657)<br>U.S. 1999-2019 (n=492)<br>Global 2007-2016 (n=61) | Alberta: 274 human, 84 cattle<br>U.S.: 312 human, 38 cattle<br>Global: 39 human, 22 cattle | Structured coalescent with 3 demes (1 per geography); $N_e$ initial value 10 and Weibull distribution (shape=1, scale=100); migration initial value 1.0 and Exponential distribution (mean=1) | HKY with empirical frequencies and discrete Gamma site model with 4 categories | Relaxed log-normal with initial value $1.5 \times 10^{-4}$ and Log-Normal distribution ($M=1.5 \times 10^{-4}$, $S=1.5$, with mean in real space) |

<sup>a</sup>Primary analysis was run using four different random seeds to confirm that all converged to the same solution. The four runs were combined to produce the final maximum clade credibility tree and state transition estimates. Model priors from this analysis were also used in an analysis in which draws were taken from the prior distribution, as opposed to the posterior distribution, to confirm that the final results were not overly influenced by the choice of priors.
<sup>b</sup>Clade A isolates were excluded from these analyses given the very small number available from any locale and clade A's divergence from the rest of the clades.

**Supplemental Table S2.** Estimated migrations from structured coalescent analysis of Alberta, U.S., and global isolates, excluding clade A.

| Migration Direction | Mean | 95% HPD |
| --- | --- | --- |
| Alberta to Alberta | 589 | 570, 609 |
| Alberta to Global | 8 | 4, 11 |
| Alberta to U.S. | 12 | 6, 16 |
| Global to Alberta | 68 | 57, 78 |
| Global to Global | 442 | 363, 511 |
| Global to U.S. | 296 | 267, 335 |
| U.S. to Alberta | 5 | 0, 14 |
| U.S. to Global | 13 | 0, 35 |
| U.S. to U.S. | 104 | 10, 184 |

Abbreviation: HPD, highest posterior density interval

